## Supplementary figures and descriptions of movie files for "Correlative single-molecule and structured illumination microscopy of fast dynamics at the plasma membrane"

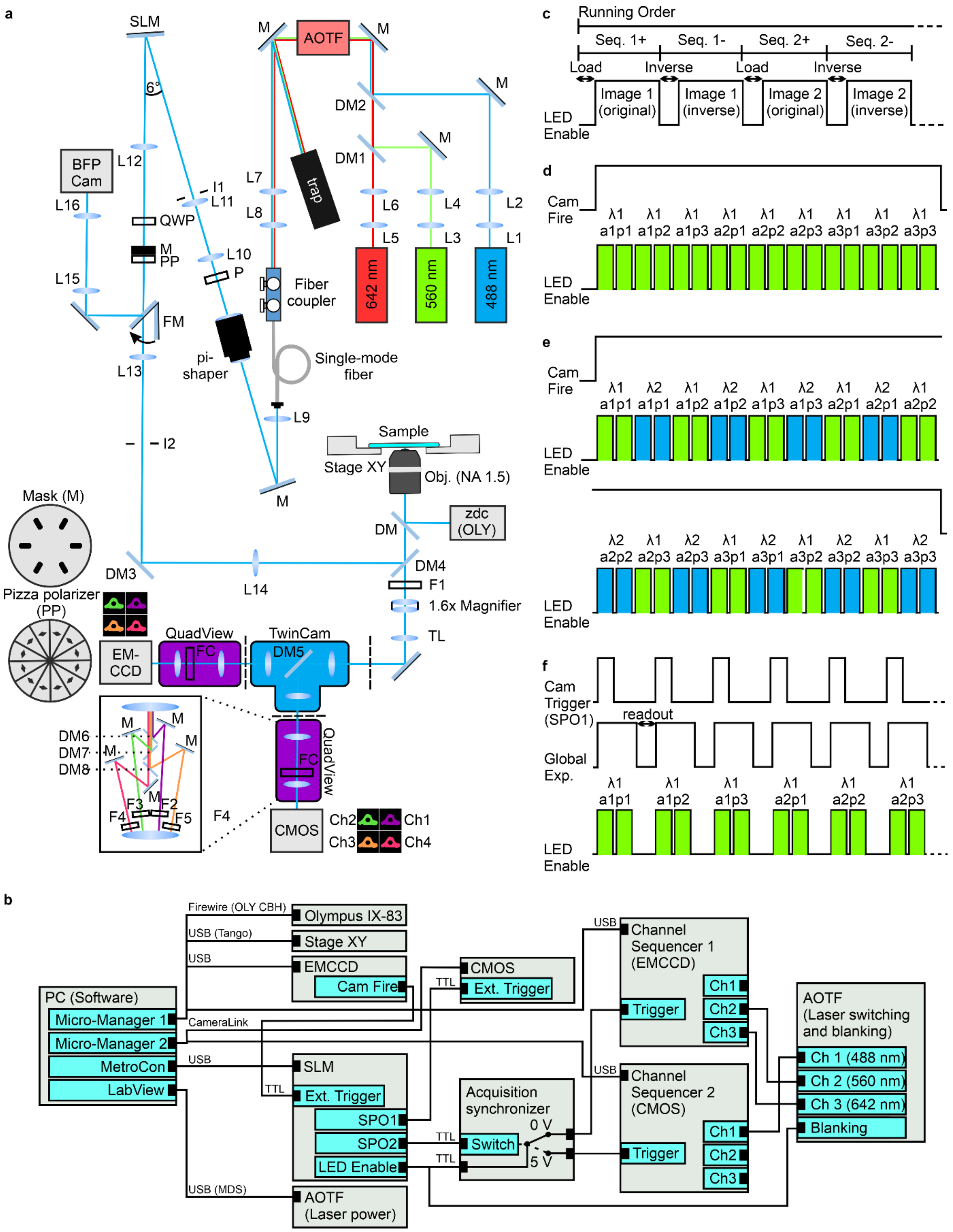


**Supplementary Fig. S1 Microscope and hardware synchronization design for (combined) single-molecule imaging and super-resolution SIM. a** Technical drawing of microscope setup (details in method section). L#: lens, M: mirror, AOTF: acousto-optical tunable filter, I: iris, P: polarizer, FM: flip mirror, SLM: spatial light modulator, QWP: quarter-wave plate: M: mask, PP: pizza polarizer, DM: dichroic/polychroic mirror, F#: bandpass filter, TL: tube lens: FC: filter cube, Ch#: fluorescence channel. **b** Diagram of software control and hardware-based fast triggering for combined single-molecule imaging using EMCCD and SIM using CMOS camera. (details in method section). Connections shown allow for 2-color SMLM (560 nm & 642 nm) and 1-color SIM (488 nm). Other configurations require reprogramming of the two channel sequencers and correct physical connection with the AOTF channels. The acquisition synchronizer is controlled by the SLM (SPO2) to send correct trigger signals (LED enable) to the sequencers. **c** Diagram showing architecture of running orders programmed on the SLM. A running order links a sequence file (timing of image display) with a binary image to be displayed. Each image needs to be inverted and displayed again to guarantee DC balancing. The LED enable TTL signal was used for laser blanking and as laser channel sequence trigger. **d** Diagram for 1-color SMLM imaging with homogenous illumination. Nine different binary images are sequentially loaded to the SLM during a single exposure of the EMCCD camera (Cam Fire signal). Each image corresponds to a specific wavelength (λ#, lambda), pattern orientation (a#, angle) and pattern phase (p#, phase). **e** Diagram for simultaneous 2-color SMLM for live-cell single-molecule tracking. Three-color SMLM followed the same principle. **f** SIM required external triggering of CMOS camera to synchronize global exposure with laser illumination and correct image display on SLM. Diagram shows acquisition of first six images for 1-color SIM. Multi-color SIM was realized by acquisition of nine SIM frames for each color one after another starting with longest wavelength.

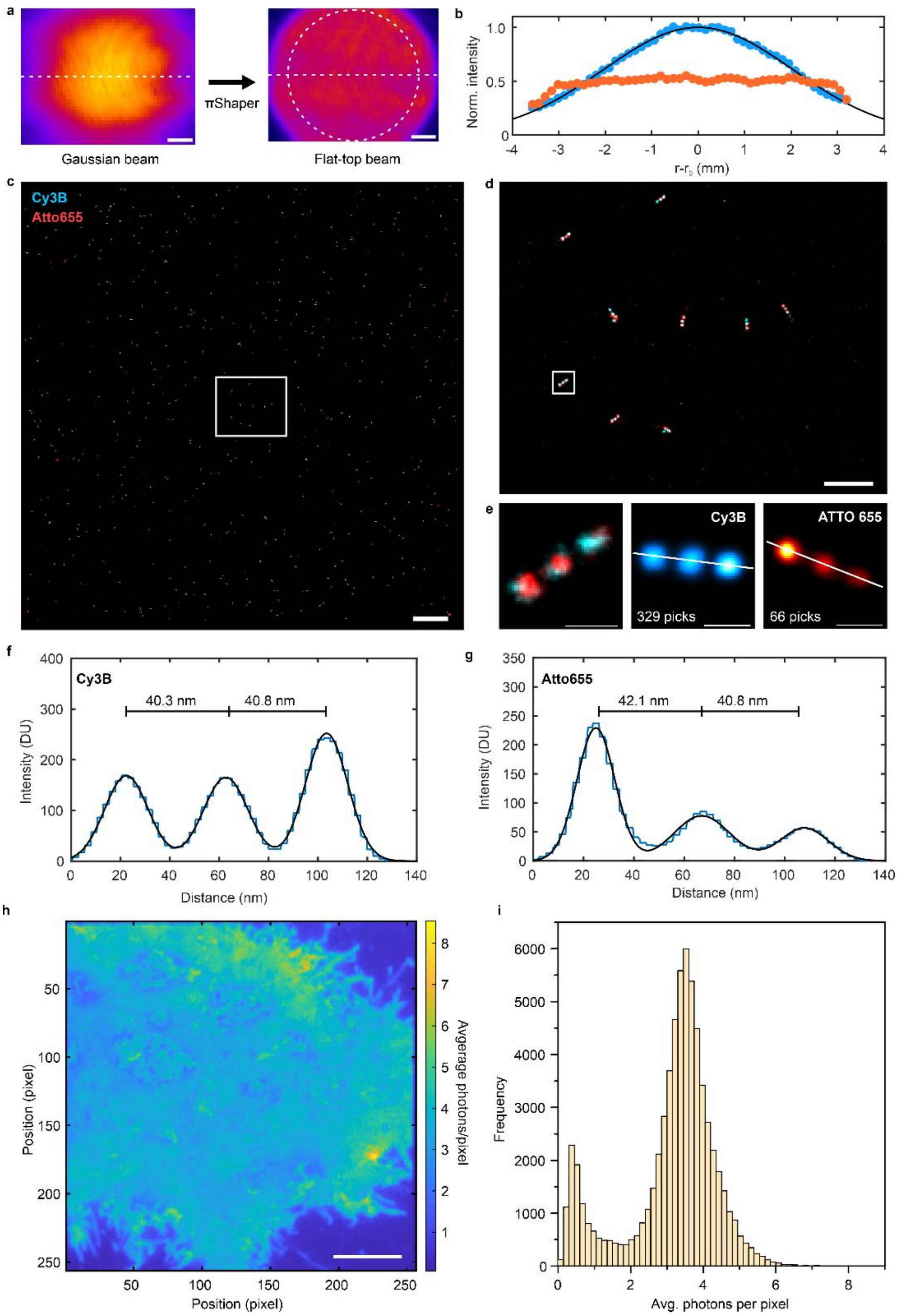


**Supplementary Fig. S2 Beam shaping performance and homogenous TIR illumination for single-molecule super-resolution and stained plasma membrane in live cells**. **a** Intensity profile before and after πShaper measured by inspection camera. Scale bar: 1 mm. Dashed circle indicates beam area illuminating SLM. **b** Line profiles along dashed lines of beam profiles in **a**. Blue circles: Gaussian profile of input beam. Black line: Gaussian fit. Orange profile illustrates clear flat-top profile from πShaper‘s output. **c-g** Dual-color DNA PAINT of DNA origamis (Gattaquant Nanorulers PAINT 40RY). **c** Overview image and zoom into highlighted region **d** as well as a single nanoruler and averaged structures for the Cy3B and ATTO 655 channels **e**. Number of picked structures for averaging are indicated. **f, g** Intensity cross-sections from averaged nanorulers in **e**. Indicated mean distances of each target on the origami was determined by Gaussian fitting using three populations. **h, i** TIRF imaging of HeLa cells expressing farnesyl-GFP for labeling of the plasma membrane. **h** Average intensity projection from 200 frames. Scale bar: 5 µm. **i** Corresponding pixel intensity histogram.

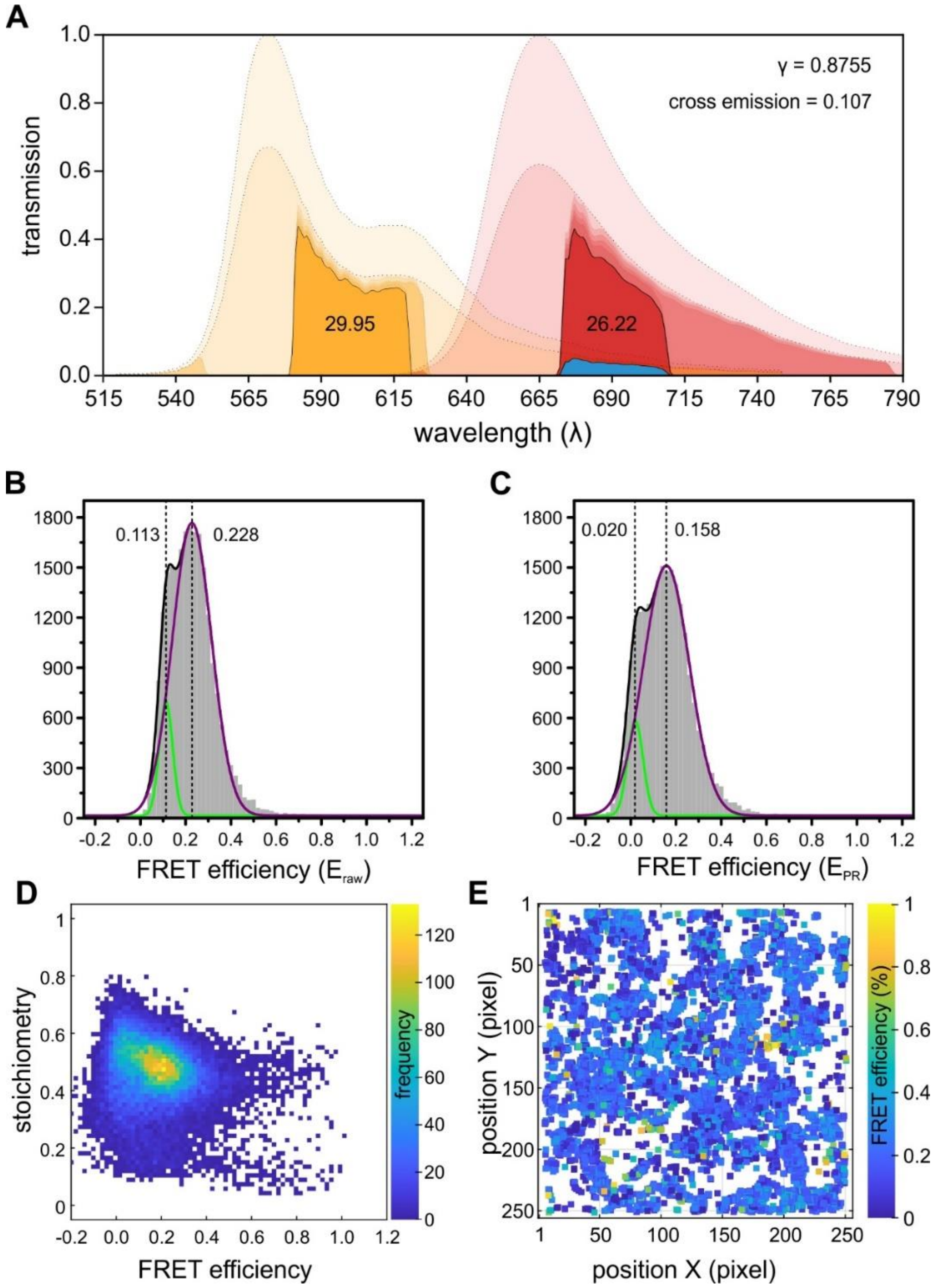


**Supplementary Fig. S3 Quantifying smFRET efficiencies. a** Calculation of the correction parameters for determining smFRET efficiencies taking the emission spectra of donor (orange) and acceptor (red) into account as well as the transmission characteristics of the emission filters. **b, c** FRET efficiency histogram lacking correction of cross-emission from the donor into the acceptor channel **b** and cross-emission corrected proximity ratio $E_{PR}$ **c**. **d** Plot of the smFRET efficiency vs. the complex stoichiometry for all co-localized donor and acceptor signals. **e** Pooled FRET efficiency map of individual TpoR dimers diffusing in the plasma membrane showing homogeneity in space and time. Histograms were fitted with a bimodal Gaussian.

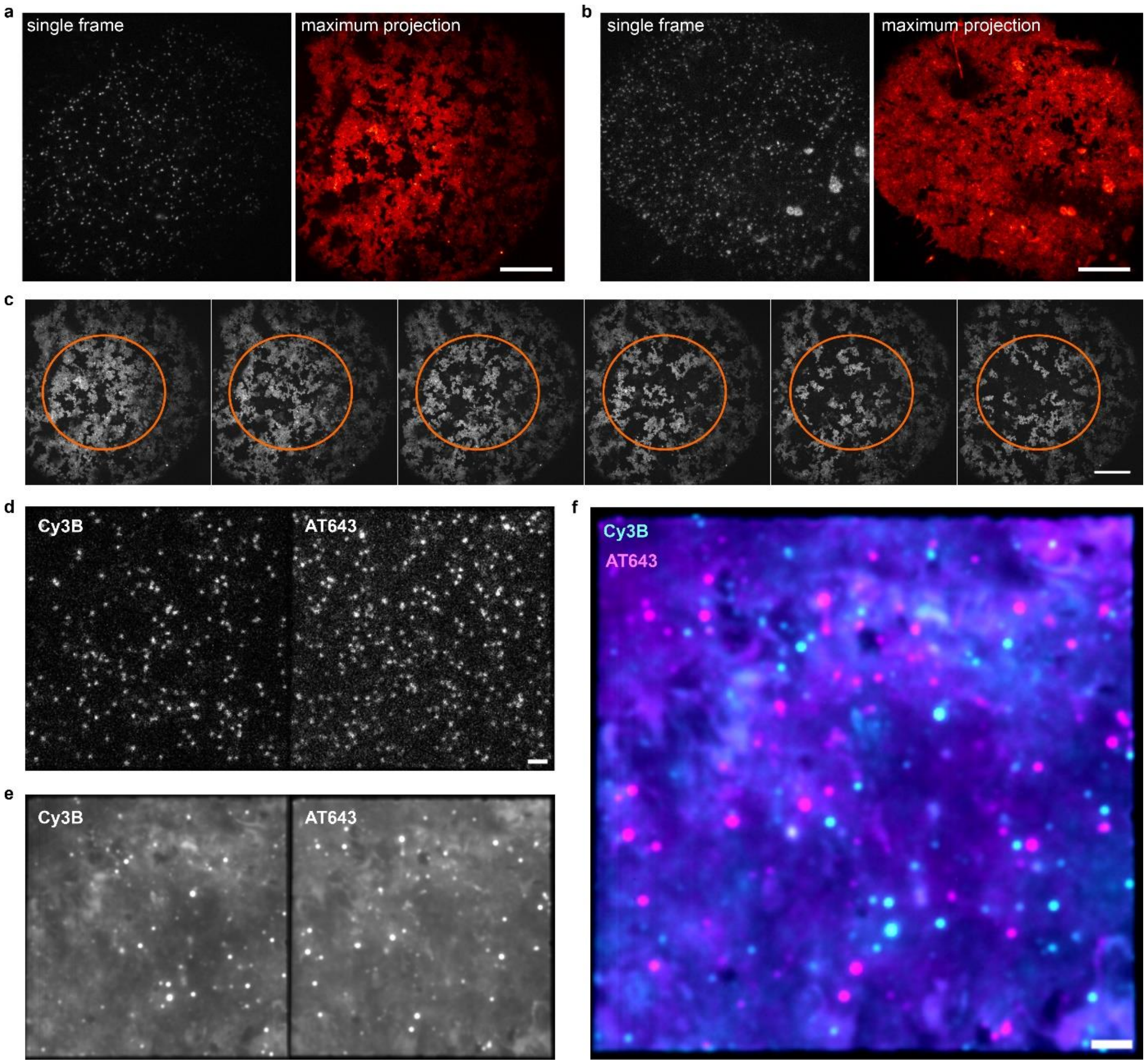


**Supplementary Fig. S4 Long-term single and dual color tracking and localization microscopy (TALM). a, b** Single-molecule imaging of mXFP-TpoR labeled with $^{AT643}$EN upon Gaussian **a** and flat-top illumination **b**. A single frame (left) and a maximum intensity projection from 1,000 consecutive frames (right) are shown in each panel. **c** Biased photobleaching upon illumination with a Gaussian beam profile, images sequentially show a maximum projection of 500 consecutive frames, of total 3,000 frames. **d-f** Dual-color TALM at high molecule density. Single frames **d** and average intensity projection of 9,000 consecutive frames **e**. **f** Overlay of the average intensity projections of both channels shown in **e**. Scale bar: 2 µm.

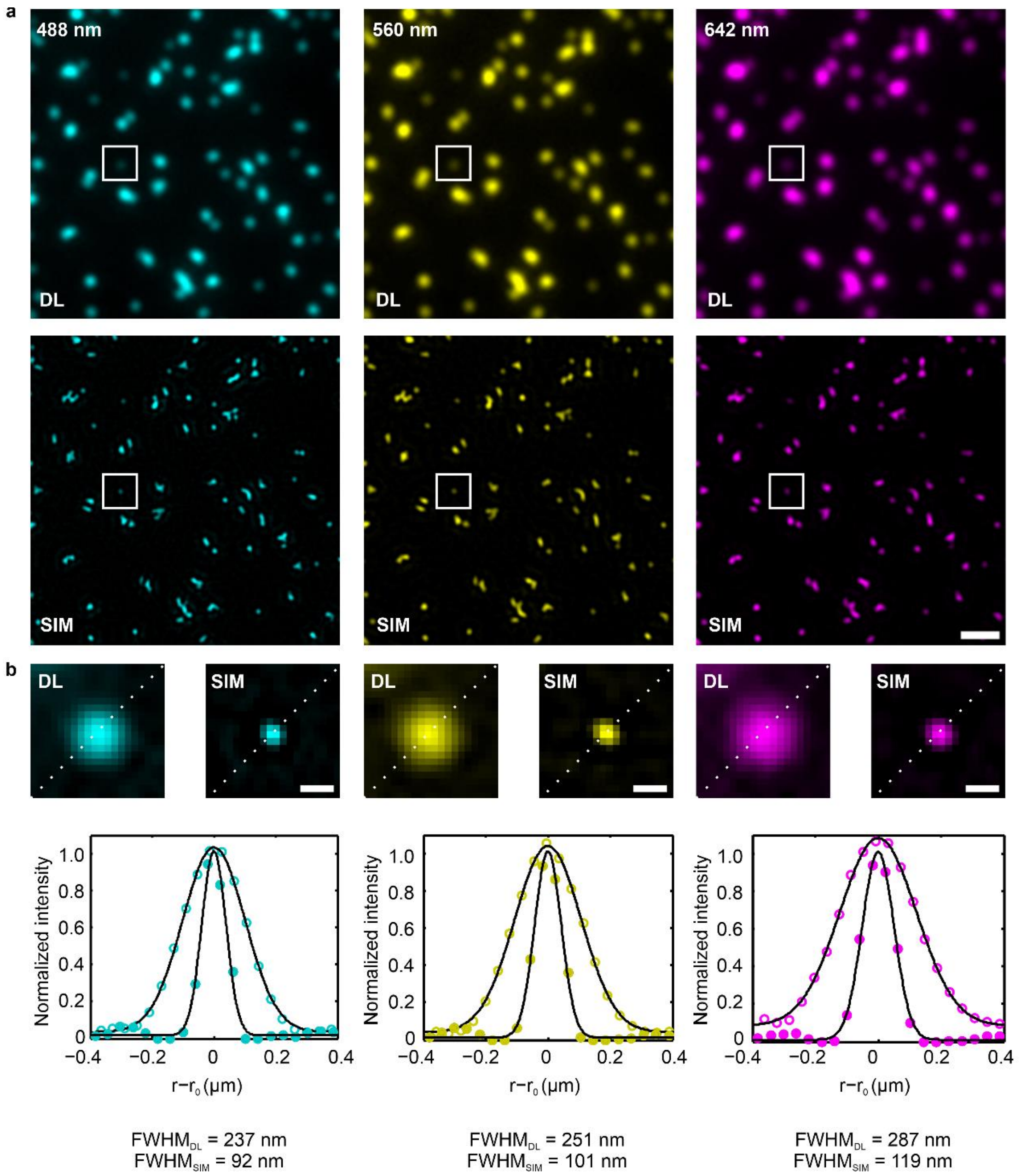


**Supplementary Fig. S5 Three-color SIM imaging of fluorescent nanoparticles. a** Diffraction-limited (DL) images (top) and deconvolved SIM images (bottom) of 100 nm TetraSpec$^{TM}$ microspheres in the three channels. DL images are synthetic images derived from averaging 9 SIM images. Scale bar: 1 µm. **b** Top: Zoomed image from ROI highlighted in **a** showing an individual nanoparticle by diffraction-limited images (DL) and after SIM deconvolution. Scale bar: 200 nm. Bottom: cross-sections and resolution determined as the FWHM (Gaussian fit) of the intensity distributions for all three channels. Open circles: DL cross-sections, closed circles: cross-sections from SIM reconstructions. Black lines: Gaussian fits.

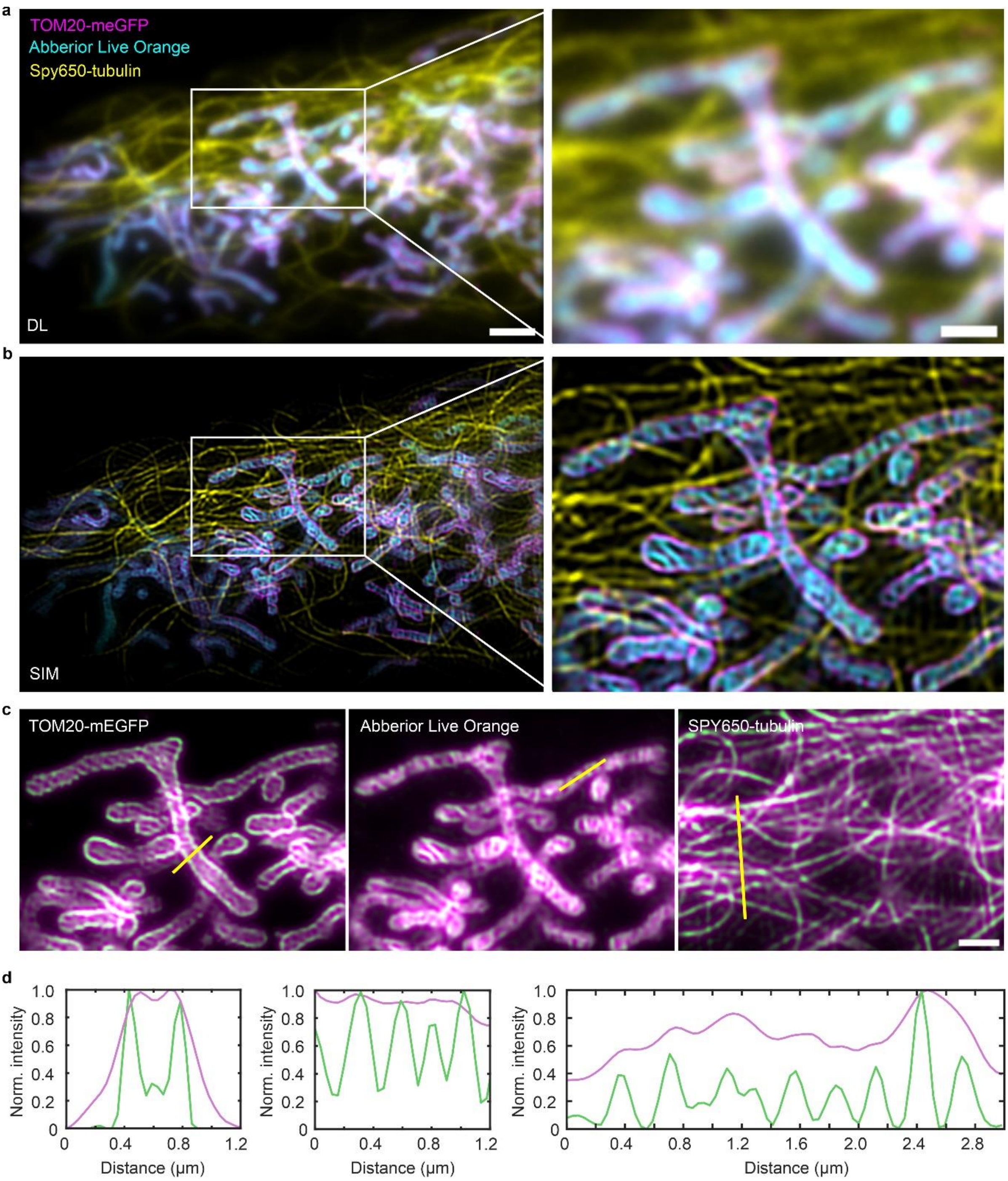


**Supplementary Fig. S6 Fast three-color GI-SIM of HeLa-cells expressing TOM20-meGFP and labeled with Abberior Mito Orange and SPY650-tubulin. a, b** Diffraction-limited (**a**) and SIM-deconvolved (**b**) three-color images showing an overview (left) and a zoomed ROI (right) highlighted by a white rectangle. Scale bars: 2 μm (left images) and 1 μm (right images). **c, d** Individual channels from right images of **a** (magenta) and **b** (green) and intensity cross-sections (**d**) along the indicated yellow lines in **c**.

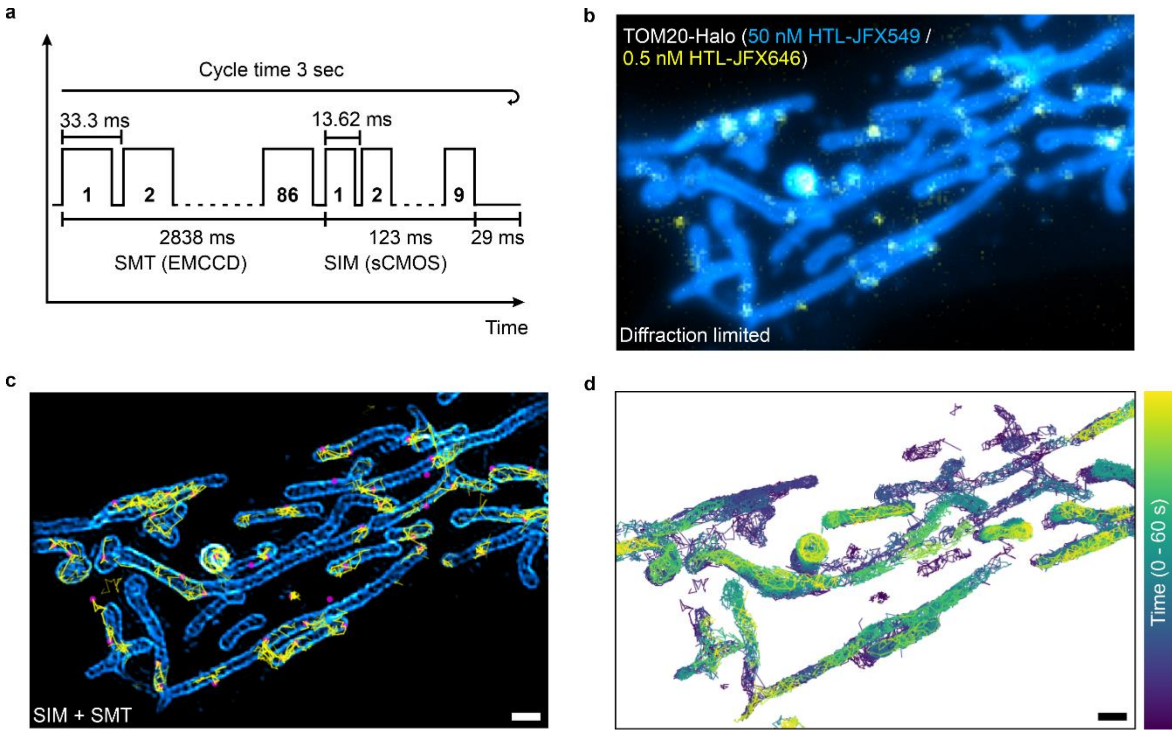


**Supplementary Fig. S7 Combination of GI-SIM and SMT imaging in live mitochondria.** **a** Diagram showing acquisition cycle of single-molecule imaging (86 frames) at 30 frames per second and subsequent acquisition of 9 SIM images with a total cycle time of 3 seconds. **b, c** Simultaneous SIM and SMT of TOM20-HaloTag in live cells by bulk labeling using 50 nM HTL-JFX549 (SIM channel) and substoichiometric labeling at 0.5 nM using HTL-JFX646 (SMT channel). Diffraction limited representation (**b**) versus processed images (**c**) using SIM reconstruction (cyan) and single-molecule localization and tracking. Magenta dots: single localizations, yellow lines: trajectories. **d** Color-coded trajectories of 5,400 consecutive frames. Scale bar: 1 µm.

### Supplementary Movies

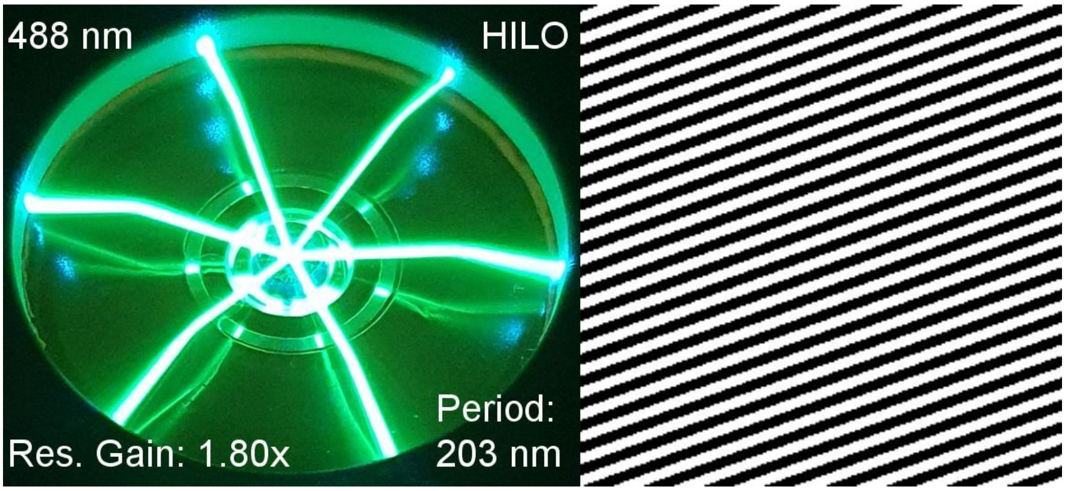


**Supplementary Movie S1**. Illustration of binary phase grating switching within single camera exposures and different illumination conditions. Here, we were illuminating a dye solution (volume: 1 ml) containing fluorescein, Texas Red and ATTO 655 at approx. 1-2 µM in PBS. Left side is showing pictures of our open custom-build sample chamber on the microscope stage. Depending on the grating period, laser is entering dye solution (HILO), propagating horizontally with respect to cover slip (grazing incidence at critical angle) or is totally reflected (TIR). Movie is showing all three channels (488 nm, 560 nm, and 642 nm) and referring to theoretical resolution gain factor for 2D-SIM as well the period of structured illumination in the focal plane of the objective.

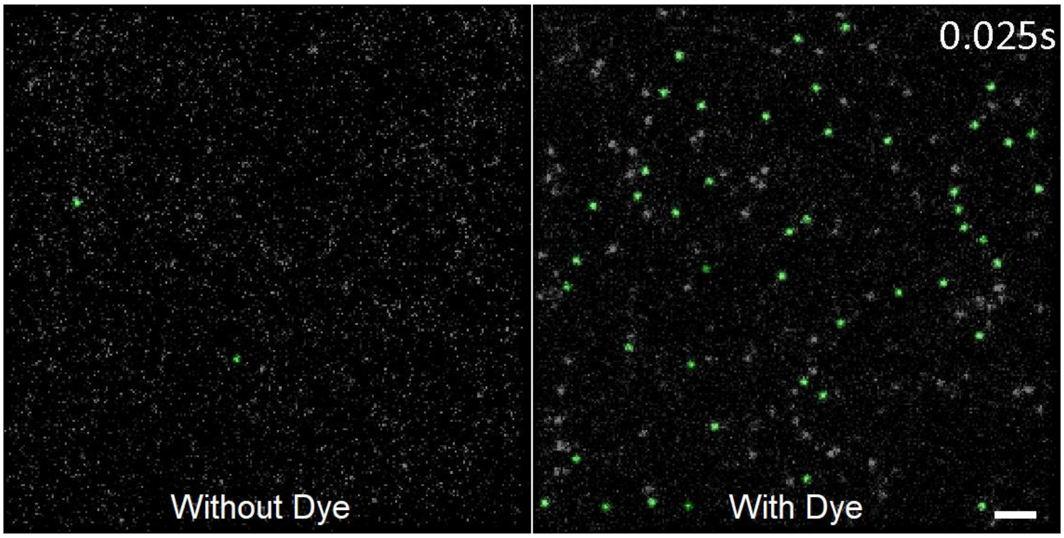


**Supplementary Movie S2**. PAINTing of immobilized reHaloTags by HaloTag ligands conjugated to MaP555 (HTL-MaP555). Tris-NTA surface loaded with reHaloTag in the absence (left) and presence (right) of HTL-MaP555. Tracked localizations of immobile trajectories longer than 2 frames are highlighted in green. Scale bar: 2 µm.

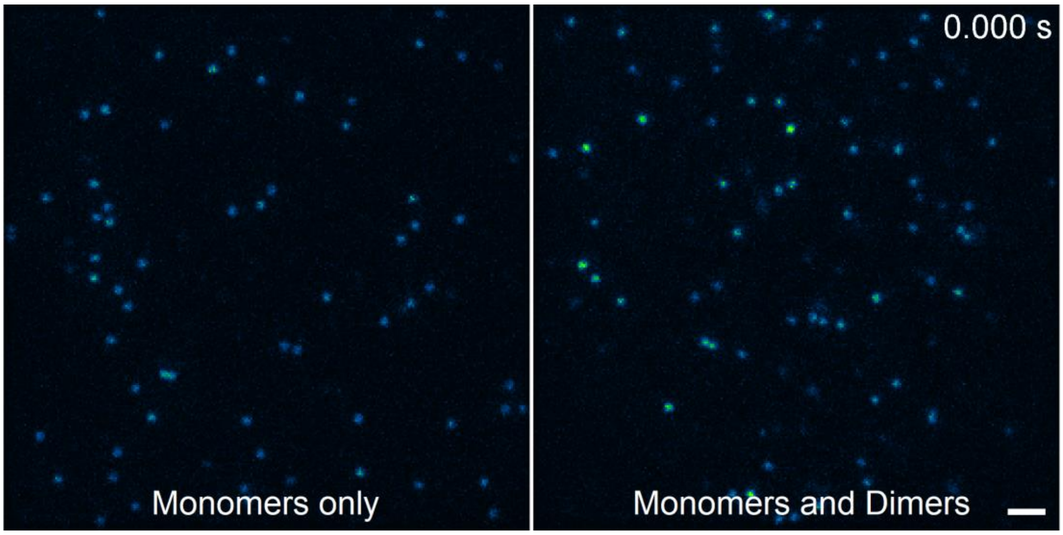


**Supplementary Movie S3**. Receptor stoichiometry by intensity analysis. ALFA-mXFP-TpoR labeled with $^{AT643}$EN in the absence (left) and presence (right) of dimerizer tdALFAnb. Scale bar: 2 µm.

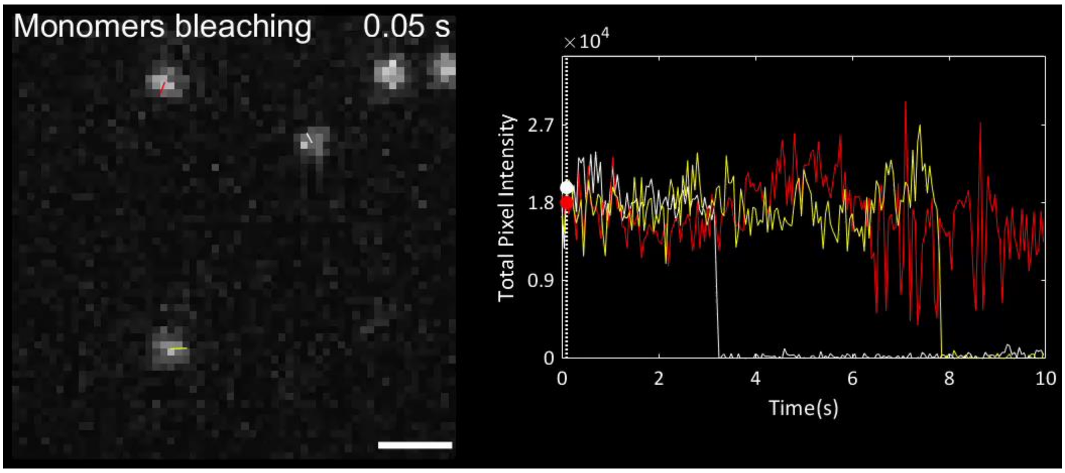


**Supplementary Movie S4**. Photobleaching of monomeric $^{AT643}$EN-labeled ALFA-mXFP-TpoR in the absence of tdALFAnb. Scale bar: 1 µm.

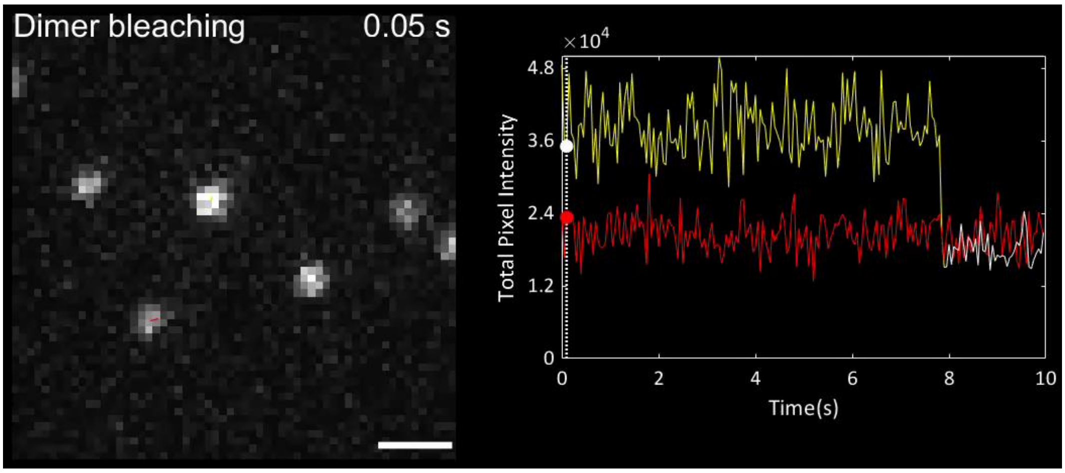


**Supplementary Movie S5**. Photobleaching of dimeric $^{AT643}$EN-labeled ALFA-mXFP-TpoR in the presence of tdALFAnb. Scale bar: 1 µm.

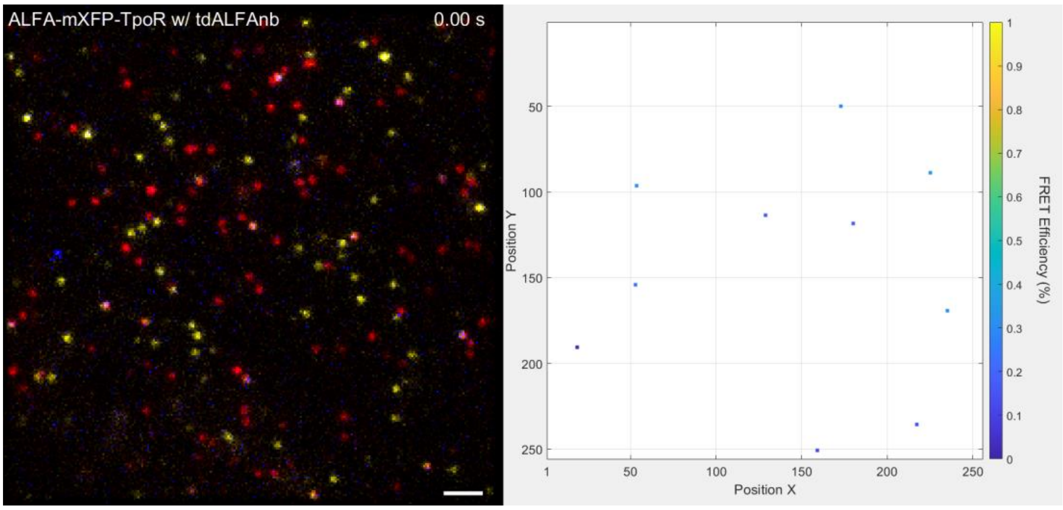


**Supplementary Movie S6**. Single-molecule ALEX-FRET detects TpoR dimers in the plasma membrane of live cells. Left: Overlay of raw data channels: Donor (yellow), acceptor (red) and FRET (blue). Single dimers are clearly visible as diffusing white signals. Scale bar: 2 µm. Right: Frame-by-frame detected FRET signals color-coded with its FRET efficiency.

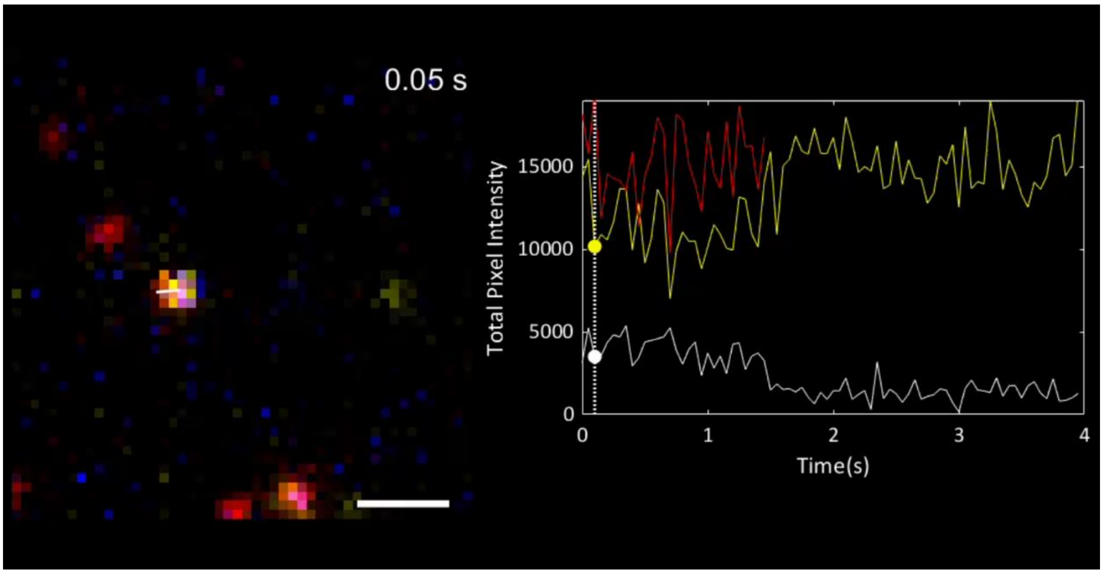


**Supplementary Movie S7**. Co-localization and co-tracking of donor (yellow), acceptor (red) and FRET channel (blue) on single complex level showing an acceptor bleaching event. Scale bar: 1 µm.

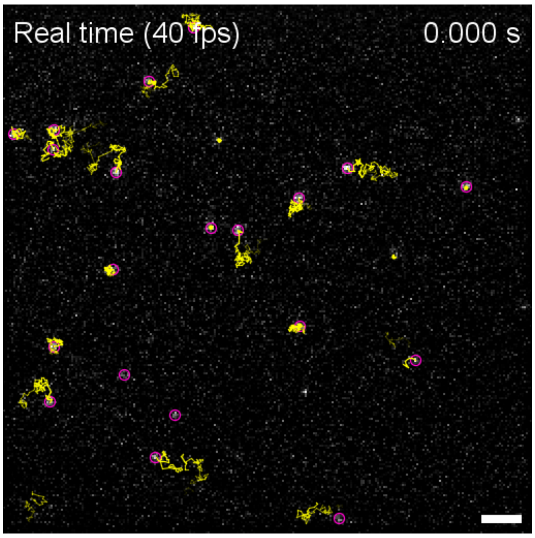


**Supplementary Movie S8**. Long-term single-molecule tracking at low receptor density. TpoR N-terminally fused to a non-fluorescent monomeric EGFP (mXFP) was expressed in HeLa cells and labeled with anti-GFP nanobodies conjugated to ATTO 643. A typical cell was imaged at 40 frames per second for 625 s (25,000 frames). Scale bar: 2 µm.

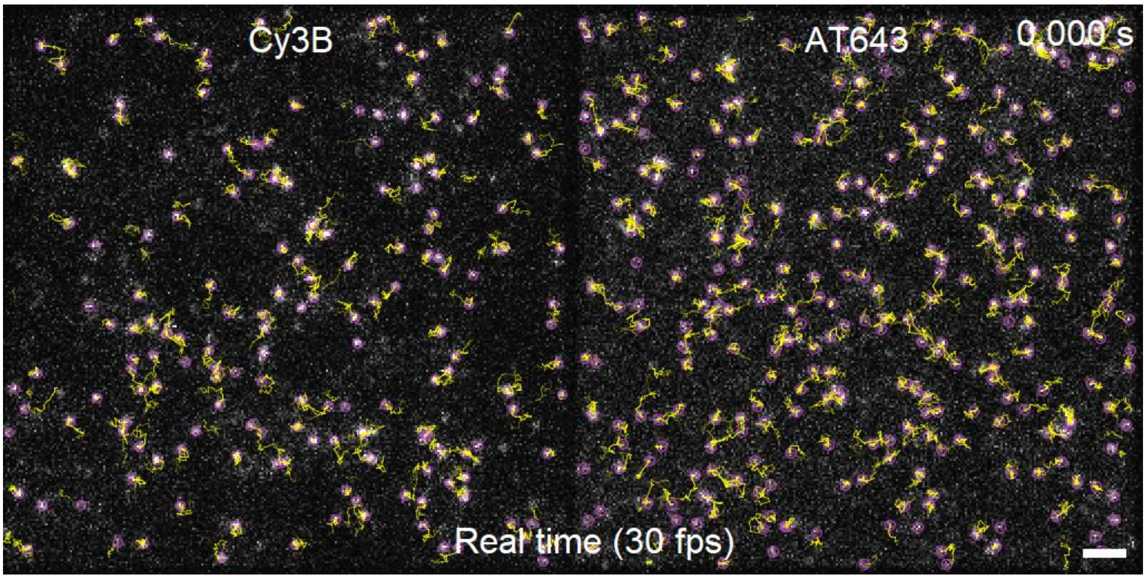


**Supplementary Movie S9**. Long-term dual-color single-molecule tracking at high receptor density. TpoR N-terminally fused to a non-fluorescent monomeric EGFP (mXFP) was expressed in HeLa cells and labeled at equimolar concentrations with anti-GFP nanobodies conjugated to Cy3B and ATTO 643, respectively. Simultaneous dual-color single-molecule tracking at 30 frames per second for 300 s (10,000 frames). Scale bar: 2 µm.

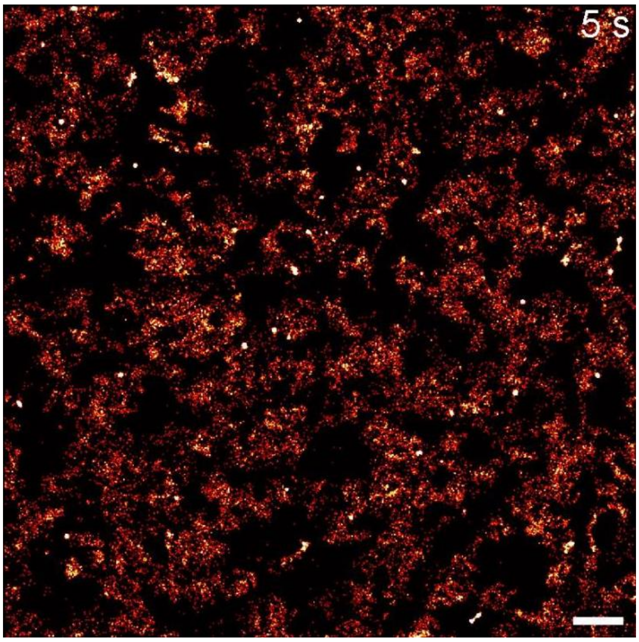


**Supplementary Movie S10**. Time-lapse localization map reveals receptor accessibility and endosome dynamics. TpoR N-terminally fused to a non-fluorescent monomeric EGFP (mXFP) was expressed in HeLa cells and labeled with anti-GFP nanobodies conjugated to ATTO 643. Single-molecule localizations of 150 consecutive frames (5 s) were binned and rendered with a moving window based on a time interval of 30 frames (1 s). Scale bar: 2 µm.

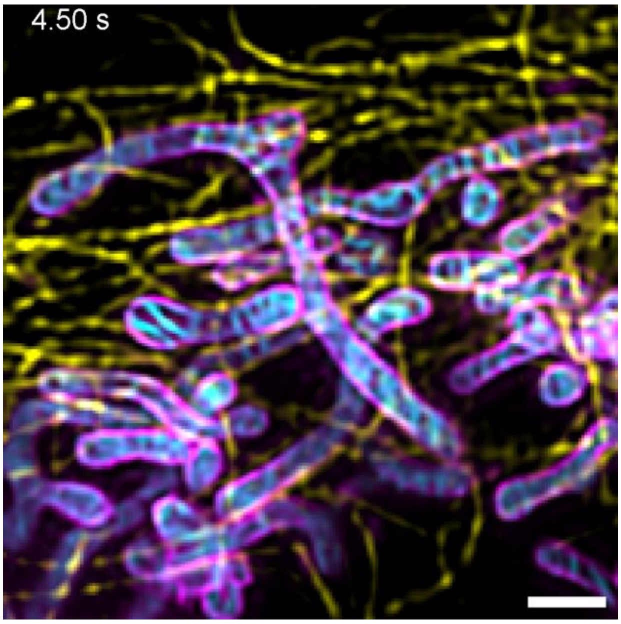


**Supplementary Movie S11**. Three-color SIM imaging of mitochondrial membrane dynamics within the microtubule network. GI-SIM of HeLa-cells expressing TOM20-meGFP (magenta) and labeled with Abberior Mito Orange (cyan) and SPY650-tubulin (yellow). Scale bar: 1 µm.

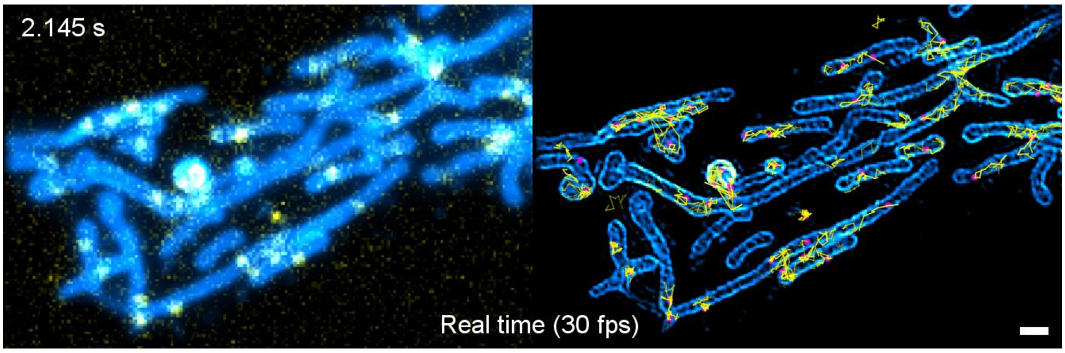


**Supplementary Movie S12**. Simultaneous SMT and SIM imaging of TOM20-HaloTag in live cells. Bulk labeling using 50 nM HTL-JFX549 (SIM channel) and substoichiometric labeling at 0.5 nM by HTL-JFX646 (SMT channel). By cycling between real-time single-molecule imaging at 30 Hz and short SIM acquisitions every three seconds, we achieved quasi-simultaneous SIM and SMT imaging in live cells. Diffraction limited representation (left) versus processed images showing SIM reconstruction (cyan) and single-molecule localization (magenta) and tracking (yellow) (right). Scale bar: 1 µm.

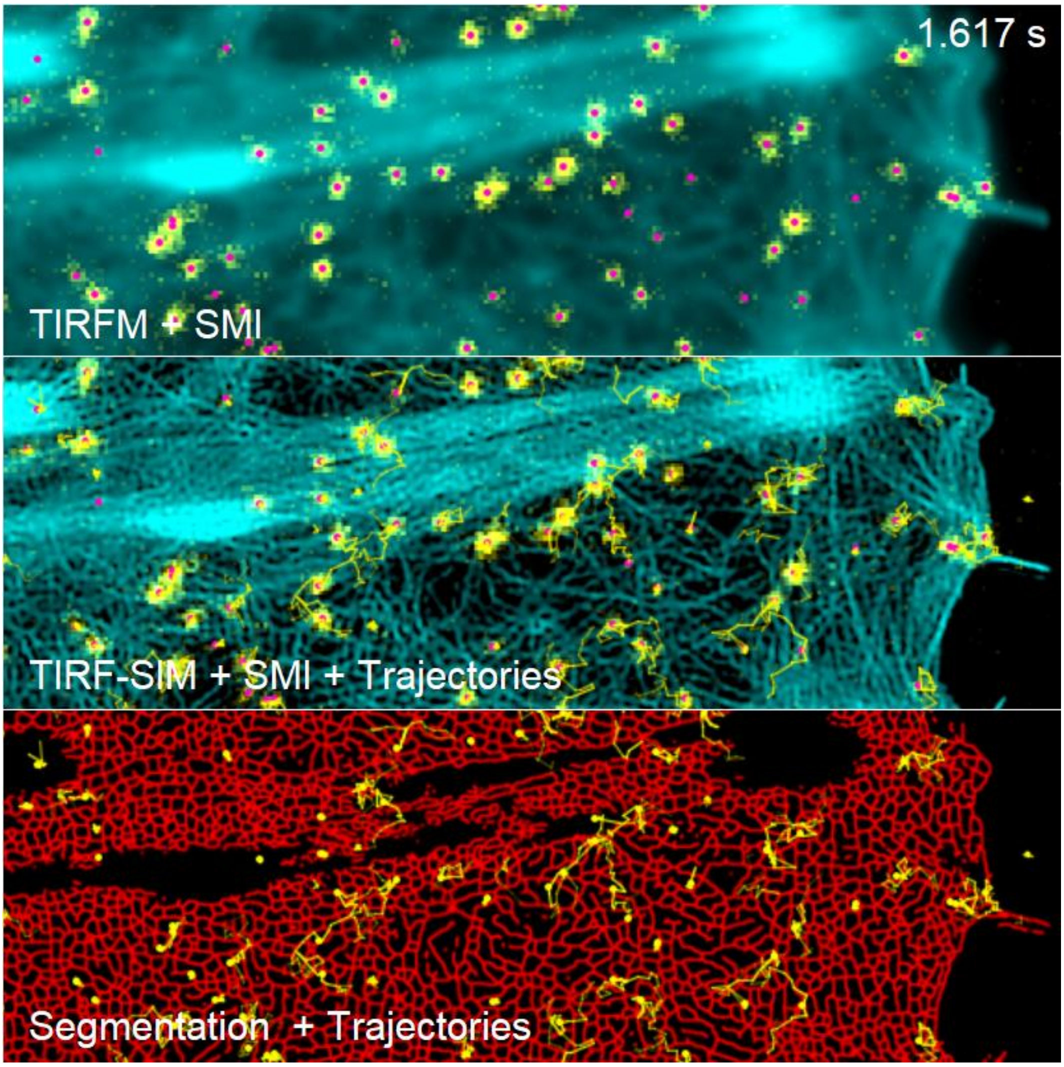


**Supplementary Movie S13**. Dynamics of the cortical cytoskeleton and Tpo receptors in the plasma membrane captured by TIRF structured illumination microscopy. TpoR N-terminally fused to a non-fluorescent monomeric EGFP (mXFP) was expressed in HeLa cells and labeled with anti-GFP nanobodies conjugated to ATTO 643. Actin cytoskeleton was imaged by LifeAct-HaloTag/JFX549. Cycling between real-time single-molecule imaging at 30 Hz and short SIM acquisitions every three seconds.  Scale bar: 2 µm.

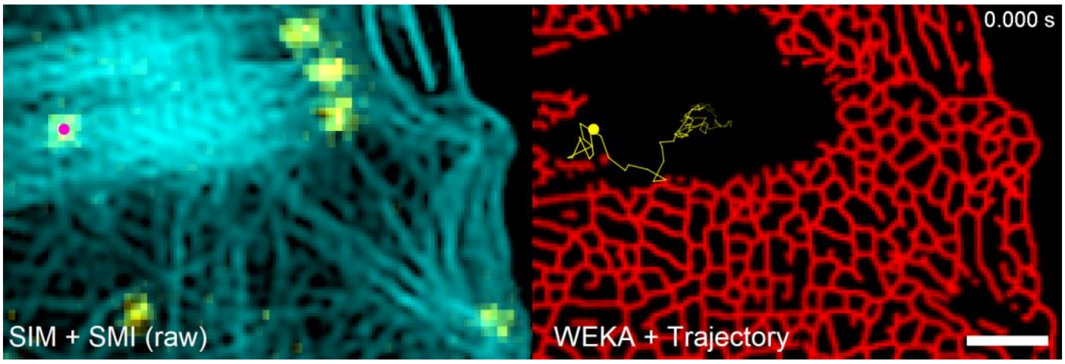


**Supplementary Movie S14**. Diffusion of TpoR in the context of clustered and corralled regions of the cortical cytoskeleton. Scale bar: 1 µm.

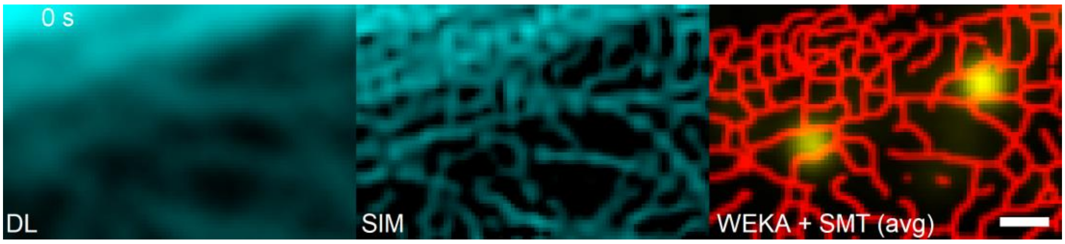


**Supplementary Movie S15**. Cytoskeletal dynamics during endocytosis of TpoR. Scale bar: 500 nm.
